# A screen for gene regulators encoded by Kaposi’s sarcoma-associated herpesvirus uncovers a novel role for the ORF42 protein in regulating the production of cellular and viral proteins

**DOI:** 10.1101/375386

**Authors:** Matthew Butnaru, Hembly Rivas, Marta M. Gaglia

## Abstract

The tight control of viral and host gene expression is critical to the replication of herpesviruses, including the gamma-herpesvirus Kaposi’s sarcoma-associated herpesvirus (KSHV). While some of the KSHV proteins that contribute to viral and host gene regulation are known, it is clear that there are additional uncharacterized contributing viral factors. Identifying these proteins and their role in gene regulation is important to determine the mechanistic underpinnings of the complex replication cycle of KSHV. Through a reporter-based screen, we have identified several new potential KSHV-encoded gene regulators, including the previously uncharacterized protein ORF42, which we find stimulates global protein production upon overexpression. We have generated an ORF42-null virus, which revealed that ORF42 is required for wild-type levels of virus production. Moreover, global protein synthesis and the accumulation of viral proteins are reduced in infected cells in the absence of ORF42, suggesting that ORF42 regulates protein synthesis during infection. A comparison of the effects of ORF42 on the levels of RNA and protein suggests that ORF42 acts post-transcriptionally to control protein levels. In addition to gene regulation, ORF42 may have other functions in virion formation, as it is found in viral particles, which is consistent with the described roles of the ORF42 homologs in alpha- and beta-herpesviruses.

**Importance:** Kaposi’s sarcoma-associated herpesvirus (KSHV) causes Kaposi’s sarcoma, an AIDS-associated malignancy that remains one of the leading causes of cancer deaths in sub-Saharan Africa. Replication of the virus is important for tumor formation and inhibition of viral replication may be used for treatment. The correct levels and temporal expression of viral and host genes during KSHV replication are key to viral replication, but the mechanisms that control this regulation remain enigmatic. Here we identify several new KSHV proteins involved in viral and cellular gene regulation and characterize the previously unstudied KSHV ORF42 protein in regulation of viral and host protein levels and efficient formation of viral progeny.

## Introduction

Kaposi’s sarcoma-associated herpesvirus (KSHV) is a gamma-herpesvirus and the etiological agent of Kaposi’s sarcoma (KS), as well as two rare lymphoproliferative diseases (primary effusion lymphoma and the B cell variant of multicentric Castleman’s disease) and KS-associated herpesvirus inflammatory cytokine syndrome (1). KS predominantly arises in individuals with compromised immune responses, such as AIDS and transplant patients, although it was also endemic in Sub-Saharan Africa prior to the AIDS epidemic and remains very prevalent in this area. KSHV establishes a long-term latent infection in patients, with periodic reactivation of the lytic cycle, which results in virus production. Both latent infection and lytic reactivation are important for KS tumorigenesis. In particular, treatment of KS patients with ganciclovir, an inhibitor of herpesvirus lytic replication, causes tumor regression (2), pointing to a role for lytic replication in the disease. Although these findings suggest that targeting lytic replication could reduce tumor formation, many aspects of the lytic cycle of KSHV are poorly characterized, including the function of many viral proteins, which limits the ability to develop approaches to reduce replication.

The control of both host and viral gene expression is key for the successful completion of the replication cycle of KSHV. Transcriptomic analysis of KSHV-infected cells has revealed dramatic changes in host mRNA levels during lytic infection, which presumably contribute to promoting viral replication and suppressing innate immune responses (3). Understanding the mechanism and viral factors that control cellular gene expression in infected cells will provide information and tools to uncover the specific role of this control in viral replication. However, at present only a few of the KSHV proteins that modulate host gene expression are known, most notably the ribonuclease (RNase) ORF37/SOX (4), and they do not fully explain the dramatic cellular gene expression changes detected in genome-wide studies.

In addition to these changes in cellular genes, the ~100 genes of KSHV are also expressed in a highly regulated fashion in relation to the progression of the replication cycle (5). During latent infection, only a few viral genes are expressed. In contrast, all the KSHV genes are transcribed during the lytic cycle, but their expression occurs in three waves: immediate early, early and late. Immediate early proteins and early proteins control viral and cellular gene expression, viral DNA replication and immune evasion. Late genes are only expressed after viral DNA replication and generally encode proteins involved in virion formation and egress. Our knowledge of the viral factors involved in the regulation of viral protein production and of the cellular mechanisms they usurp is expanding, but remains incomplete. In particular, how post-transcriptional mechanisms contribute to shaping the complex pattern of viral gene expression is largely unknown.

To uncover new KSHV factors that regulate host and viral gene expression, we carried out a reporter screen and overexpressed individual KSHV proteins. We examined expression of several reporters with different constitutive promoters and regulatory regions to identify KSHV proteins that have the capacity to broadly affect gene expression. We identified several new KSHV candidate gene regulators, and we characterized the function of one of these proteins, ORF42, in the KSHV replication cycle. KSHV ORF42, which was previously uncharacterized, is one of the core herpesviral proteins. Homologs are present in all herpesvirus subfamilies, although the sequence identity among the subfamilies is limited (less than 20%). Null mutations of the ORF42 homolog UL103 in the beta-herpesvirus human cytomegalovirus (HCMV) and the UL7 homolog in the alpha-herpesviruses pseudorabies virus (PRV) and herpes simplex virus 1 (HSV-1) result in a decrease in virus production (6–9). This attenuation is thought to arise from defects in virion formation and potentially directly in viral egress (6, 7). However, there is evidence that UL7 and UL103 may have additional biological activities. One study reported that HSV-1 UL7 may regulate early lytic gene expression during *de novo* infection (10). Similarly, proteomic analyses of HCMV UL103-associated proteins revealed interaction with several cellular antiviral proteins including the DNA sensor IFI16 (11). Hence, KSHV ORF42 may also l· have multiple activities in the viral replication cycle.

Using an ORF42-null KSHV, we demonstrate that during infection ORF42 promotes a small but ι significant increase in global protein synthesis, and is thus necessary for normal accumulation of ‘ viral proteins. In addition, loss of ORF42 attenuates viral replication, reducing the number of viral particles formed and released. Our results point to a novel function for ORF42 as a post-transcriptional regulator of KSHV and host gene expression.

## Results

### A reporter screen identifies new KSHV-encoded candidate regulators of gene expression

Dysregulation of host gene expression contributes to cellular takeover during infection and l· understanding the underlying mechanisms of host gene regulation will be helpful to define how this control contributes to viral replication. In addition, the same viral factors may shape both i cellular and viral gene expression, and thus contribute to the temporal regulation of viral genes. Given the wide-ranging changes in host gene expression during KSHV infection (3) and to cast a wide net, we designed a reporter-based screen to identify viral proteins that broadly modulate o gene expression, regardless of the protein coding region, promoter, or the 5’ and 3’ untranslated I region (UTR). Indeed, many viruses encode proteins that broadly modulate gene expression, rather than affecting a select group of proteins, including the KSHV RNase ORF37/SOX (4). We ; chose three reporters with promoters and UTRs that drive constitutive, but low level, protein expression in mammalian cells (Fig.1A): a luciferase reporter driven by the herpes simplex virus l· thymidine kinase promoter with a SV40 3’ UTR, a GFP reporter driven by the long terminal repeats of the Moloney murine leukemia virus, and a DsRed reporter driven by the human i phosphoglycerate kinase (hPGK) promoter with a bovine growth hormone 3’ UTR. The ‘ relatively low levels of constitutive expression of these reporters allowed us to detect small changes in reporter expression. Luciferase activity and GFP/DsRed protein levels in HEK293T cells were measured as readouts.

**Figure 1.**
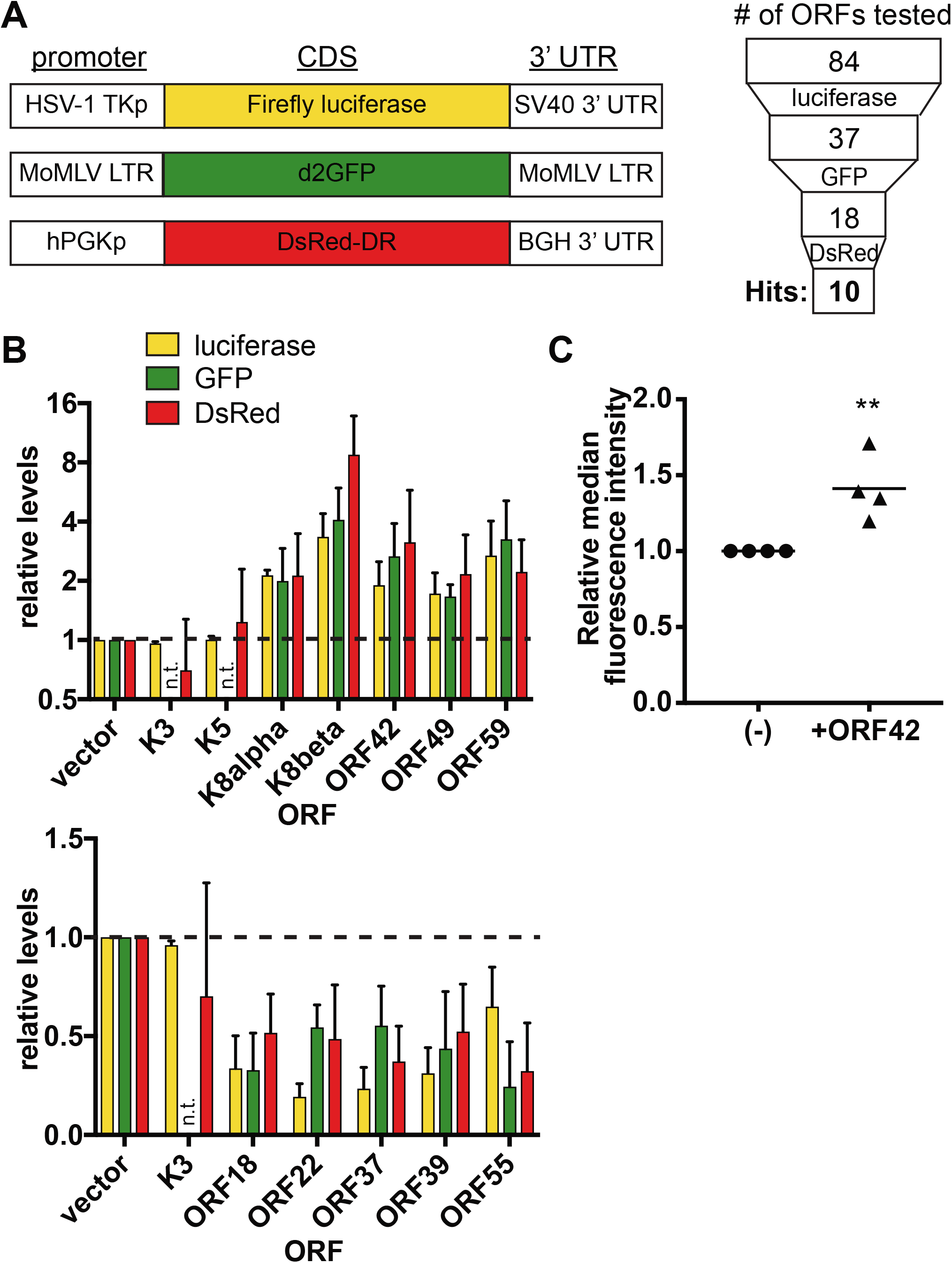
A reporter screen identifies KSHV regulators of gene expression. (A) Schematic diagram of the reporter constructs with indicated promoter, coding sequences and 3’ UTR regions *(left).* The number of viral ORFs that were tested at each stage and that passed testing is indicated in the diagram on the right. (B) The indicated C-terminally Strep-tagged KSHV ORFs were co-transfected in HEK293T with individual reporter constructs listed in (A) for 24 hrs (luciferase/GFP) or 48 hrs (DsRed). Reporter protein levels were measured by luciferase assay (luciferase) and western blot (GFP and DsRed, using tubulin or actin as loading controls). The graphs show the fold change in luciferase activity or protein levels relative to cells transfected with empty vector for ORFs that passed testing and for K3 and K5, which did not increase gene levels and are included as negative controls. ORFs that increased gene expression are shown in the graph on the top, while ORFs that decreased gene expression on the bottom.N.t. = not tested. (C) Protein synthesis rates were measured using AHA-based metabolic labeling in HEK293T cells transfected with either vector (“(-)”) or an untagged ORF42 expression construct. 24 hours after transfection, nascent proteins were labeled using AHA coupled to a fluorescent dye and fluorescence was measured using flow cytometry. The median fluorescence intensity of AHA-treated cells for four replicate experiments is plotted relative to vector-transfected cells. * = *p* < 0.05 (Student’s t-test).

To express the KSHV proteins, we used an expression library for 84 KSHV ORFs generated by the Glaunsinger lab and previously used for protein-protein interaction mapping (12). Although we did not verify expression of all the ORFs, all the constructs were shown to express the corresponding proteins in the same cell type in the Davis *et al.* study These five proteins, thus, may also function12). As detailed in (Fig. 1A, we progressively narrowed down the number of tested ORFs, at each step continuing only with those that up- or down-regulated reporter expression by at least 1.5-fold. At the end of the screen, we identified ten ORFs that consistently altered reporter protein levels. These included the known regulator of host gene expression ORF37/SOX (4) and five proteins that have previously been linked to viral gene regulation, ORF18, ORF49, ORF59 and the two K8 isoforms (Fig. 1B) (13–16). These five proteins, thus, may also function as regulators of host gene expression, because they can influence the levels of reporter proteins controlled by non-KSHV promoters and regulatory regions. In addition, we identified four proteins not previously linked to gene expression, ORF22, ORF39, ORF42, and ORF55. Of these proteins, three (ORF22, ORF39 and ORF55) reduced reporter levels and one (ORF42) increased reporter levels. While all four ORFs remain uncharacterized in KSHV, their homologs in other herpesviruses have roles in virion formation or are envelope glycoproteins (6, 7, 17–22). The results of our screen suggest that ORF22, ORF39, ORF42 and ORF55, may have additional or alternative functions in gene regulation in KSHV. We thus conclude that our multi-reporter screen identified four potential new KSHV regulators of gene expression and pointed to a wider role in host as well as viral gene expression for five additional proteins.

### KSHV ORF42 increases global nascent protein production

We decided to focus on the ORF42 protein, the only novel candidate that increased reporter expression. To confirm that the effect of ORF42 was not limited to the reporters we selected, we tested how ORF42 affected global nascent protein production. ORF42-overexpressing cells were treated with the Click-able methionine analogue L-azidohomoalanine (AHA), which is incorporated into nascent proteins. L-AHA was then conjugated to a fluorescent dye and AHA incorporation into newly synthesized proteins was measured by flow cytometry. The median fluorescence intensity of cells transfected with ORF42 was 40% higher than that of cells transfected with empty vector (Fig. 1C). This result indicates that ORF42 increases global protein synthesis, which is in line with the results of our screen.

### The putative gene regulator ORF42 is a late cytoplasmic protein that is required for wild-type levels of virus production

The ORF42 locus is predicted to encode a 31-kDa, 278-amino acid protein with homologs in all other herpesviruses. To study ORF42, we engineered a KSHV variant lacking expression of the ORF42 protein, KSHV ΔORF42, by replacing Serine 25 with a stop codon in the KSHV BAC16 clone using BAC recombineering (Fig. 2A). We chose this mutation because it does not alter the coding sequence of ORF43, which overlaps with the 5’ end of the ORF42 coding sequence (Fig. 2A). We also made a KSHV BAC16 variant that expresses a C-terminally Flag-tagged version of ORF42 from the viral genome (KSHV ORF42-Flag, (Fig. 2B). Because the transcription termination sequence and a portion of the coding region of ORF41 overlap with ORF42, we duplicated the sequences at the 3’ end of ORF42 to preserve expression of ORF41. We generated cell lines latently infected with KSHV WT, ΔORF42 or ORF42-Flag by transfecting the BACs in iSLK.RTA cells, which are uninfected epithelial cells that express a doxycycline-inducible copy of the KSHV master lytic regulator RTA (23). RTA overexpression triggered by doxycycline treatment leads to induction of the KSHV lytic cycle. After transfection, we isolated clonal populations of cells infected with KSHV ΔORF42 and ORF42-Flag and, as a control, two separate lines infected with wild-type (WT) KSHV.

**Figure 2.**
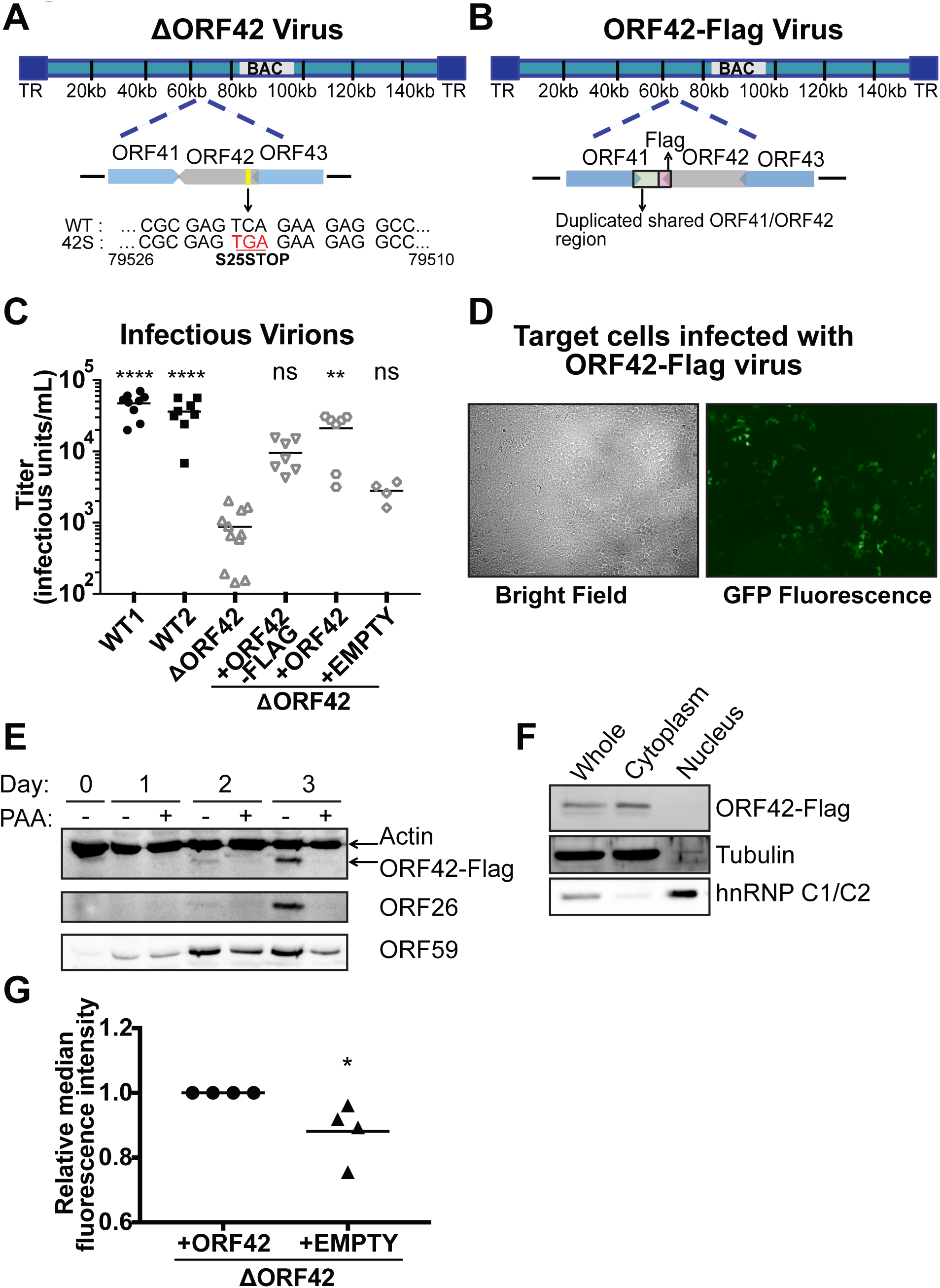
ORF42 is important for wild-type levels of infectious virus production and controls protein synthesis in infected cells. (A) Schematic diagram of KSHV ΔORF42 genome structure, including the location of nonsense mutation at serine 25. (B) Schematic diagram of the KSHV ORF42-Flag genome structure, indicating the C-terminal Flag tag position and the partial duplication of ORF41 sequences. (C) Infectious virus production from KSHV WT and ΔORF42 infected cells. The lytic cycle was induced in the indicated infected iSLK.RTA cells by addition of doxycycline (1 μg/ml) to the media. In this and other figures, the following labels are used: WT1 and WT2: two clonal lines infected with wild-type KSHV; ΔORF42: cells infected with KSHV carrying a nonsense mutation in ORF42; +ORF42-Flag: KSHV ΔORF42-infected cells transduced with C-terminally Flag-tagged ORF42; +ORF42: KSHV ΔORF42-infected cells transduced with untagged ORF42; +EMPTY: KSHV ΔORF42-infected cells transduced with empty vector. Supernatant was collected from all cell populations 6 days after induction and used to infect HEK293T target cells. Infectious units were calculated from the percentage of GFP-positive target cells measured by flow cytometry. N ≥ 4. Ns, **,**** = *p* > 0.05, or < 0.01, 0.0001, respectively, relative to KSHV ΔORF42 (ANOVA followed by Dunnet’s multiple comparison test). (D) Supernatant was collected 6 days post-induction from cells infected with KSHV carrying virus-encoded C-terminally Flag-tagged ORF42 and used to infect HEK293T target cells. Infection was assessed by fluorescence microscopy. The bright field picture shows the field of target cells, while the GFP signal identifies the infected cells. (E) Protein lysates were collected at the indicated time points from cells infected with KSHV carrying virus-encoded C-terminally Flag-tagged ORF42 and treated with the viral DNA polymerase inhibitor phosphonoacetic acid (PAA). ORF42 was detected using anti-Flag antibodies. Actin serves as a loading control, and ORF59 and ORF26 are representative early and late viral proteins, respectively. (F) Cells infected with KSHV carrying virus-encoded C-terminally Flag-tagged ORF42 were fractionated into cytoplasmic and nuclear fractions at day 4 post induction. Lysates were stained for ORF42-Flag, tubulin and hnRNP C1/C2 (cytoplasmic and nuclear loading controls, respectively). Blots in E and F are representative of at least 3 biological replicates. (G) Protein synthesis rates were measured using AHA-based metabolic labeling. The lytic cycle was induced in the indicated KSHV-infected iSLK.RTA cells by addition of doxycycline (1 μg/ml) to the media. Five days after lytic cycle induction, nascent proteins were labeled using AHA coupled to a fluorescent dye, and fluorescence was measured using flow cytometry. The median fluorescence intensity of AHA-treated cells for four replicate experiments was measured by flow cytometry and is plotted relative to KSHV ΔORF42-infected cells rescued with untagged ORF42. * = *p* < 0.05 (Student’s *t*-test).

To determine whether ORF42 is required for viral replication, we induced the lytic cycle in WT and ΔORF42 KSHV-infected cells by adding doxycycline, collected supernatant six days after induction and used it to infect naïve HEK293T cells. Because the KSHV BAC16 clone constitutively expresses GFP, the titer of infectious virions produced was measured by counting the percentage of GFP-positive infected HEK293T cells by flow cytometry. We found that cells infected with KSHV ΔORF42 produced significantly fewer infectious virions than WT-infected cells (Fig. 2C, WT = 4.7 *10^4^ ± 1.7*10^4^ infectious units/mL, ΔORF42 = 8.7 *10^2^ ± 6.4*10^2^ infectious units/mL). To make sure that this defect was due to the ORF42 mutation, we rescued ORF42 in KSHV ΔORF42-infected cells using integrated transgenes that express untagged ORF42, a Flag-tagged version of the protein, or a Flag tag-only empty vector as a negative control. Importantly, rescuing ORF42 *in trans* allowed us to test the same clonal latently infected line with and without ORF42 expression. We found that ORF42 expression in *trans* rescued virus production from the ΔORF42-infected cells, which indicates that the defect was indeed caused by the ORF42 mutation (Fig. 2C). We noticed that untagged ORF42 rescued infectious virus production more efficiently than C-terminally Flag-tagged ORF42. This result suggests that the C-terminal tag may reduce ORF42 activity, similar to what has been reported for the cytomegalovirus ORF42 homolog, UL103 (8). Despite this result, we found that the KSHV ORF42-Flag virus, which expresses ORF42-Flag from the viral genome, still produced at least some infectious virus, as shown by the appearance of GFP-positive cells after exposure to supernatant from KSHV ORF42-Flag infected cells (Fig. 2D). Nonetheless, because of the result with the ORF42-Flag rescue, we used the KSHV ORF42-Flag infected cells only to examine the spatial and temporal expression of ORF42 (Fig. 2E,F). Consistent with the annotation of the ORF42 locus, addition of the Flag tag to virus-encoded ORF42 led to the detection of a ~35-40 kDa protein (Fig 2E). Virus-encoded ORF42-Flag was expressed with the kinetics of a late lytic protein. It was not detected during latency, and it appeared at the same time as the late protein ORF26, while the early protein ORF59 was detected earlier (Fig. 2E). In addition, ORF42 production was sensitive to treatment with phosphonoacetic acid (PAA), an inhibitor of viral DNA replication, which is a hallmark of a true late protein (Fig. 2E). These data agree with results from a previous transcriptome profiling study of KSHV mRNAs in the presence of the viral DNA polymerase inhibitor cidofovir, which indicated that the ORF42 mRNA was not expressed in the absence of viral DNA replication (24). In addition, virus-encoded ORF42-Flag localized almost exclusively to the cytoplasm, as indicated by western blot analysis of fractionated cell lysates (Fig 2F). This localization was maintained throughout the lytic cycle (data not shown). Collectively, our results show that ORF42 is essential for wild-type levels of viral replication and is a late protein that localizes to the cytoplasm.

### ORF42 regulates global protein production in infected cells

In order to test whether ORF42 controls gene expression in infected cells, we again employed the AHA-based metabolic labeling assay to assess the global nascent protein production of KSHV-infected cells, specifically ΔORF42-virus infected cells rescued with WT ORF42 or empty vector. The median fluorescence intensity of KSHV lytically-infected cells was 10% lower in the absence of ORF42 (Fig. 2G). While small, this difference was reproducible and statistically significant (Fig. 2G). This result indicates that the presence of ORF42 promoted the synthesis of new proteins in infected cells. We note that it is likely that not all KSHV infected cells are lytically reactivating in this assay, and that we may thus be underestimating the effect of ORF42 on translation in infected cells. Nevertheless, this small but consistent reduction indicates that ORF42 potentiates the synthesis of a large number of proteins during infection.

### ORF42 is required for wild-type levels of late viral protein expression

Our screen had the potential to identify regulators of both cellular and KSHV gene expression, because we selected several regulatory regions of various origin (Fig. 1A). Also, the same factors can affect production of both viral and host protein, as seen for example for the ORF37/muSOX protein in the murine gamma-herpesvirus murine gammaherpesvirus 68 (MHV68) (25). To test whether ORF42 can regulate viral genes, we co-expressed ORF42 with an mRNA for KSHV ORF26, which included the coding region and the UTRs from the ORF26 genomic locus. We chose ORF26 because both ORF42 and ORF26 are late proteins, and should thus be expressed at the same time in infected cells. ORF42 co-expression led to an increase in ORF26 protein (Fig. 3A,B), but not mRNA levels (Fig. 3C). These results indicate that ORF42 can regulate viral proteins and also suggest that ORF42 regulates gene expression post-transcriptionally. We extended these results by measuring the levels of a number of viral proteins at day 4 and 6 post induction. As antibodies against KSHV proteins are limited, the tested proteins were chosen based on the availability of efficient antibodies, rather than specific functions. We observed that mutation of ORF42 reduced the levels of most of the proteins we tested, although the extent of the difference varied (Fig. 4A,B). Moreover, normal protein levels were restored upon exogenous expression of ORF42, but not by transduction of the empty vector, confirming that the defect was due to ORF42 activity (Fig. 4A,B). The decrease in protein levels was most pronounced for late proteins, such as the glycoprotein K8.1, the capsid protein ORF26 and the tegument protein ORF33 (Fig. 4B). In contrast, early proteins were only slightly decreased (Fig. 4A). This is not surprising, since ORF42 is itself a late protein (Fig. 2E), and early proteins will partially accumulate before ORF42 is expressed. Also, the alteration in late protein levels was not due to a defect in viral DNA replication, because viral DNA replication was not impaired in the ΔORF42 mutant virus (Fig. 4C).

**Figure 3.**
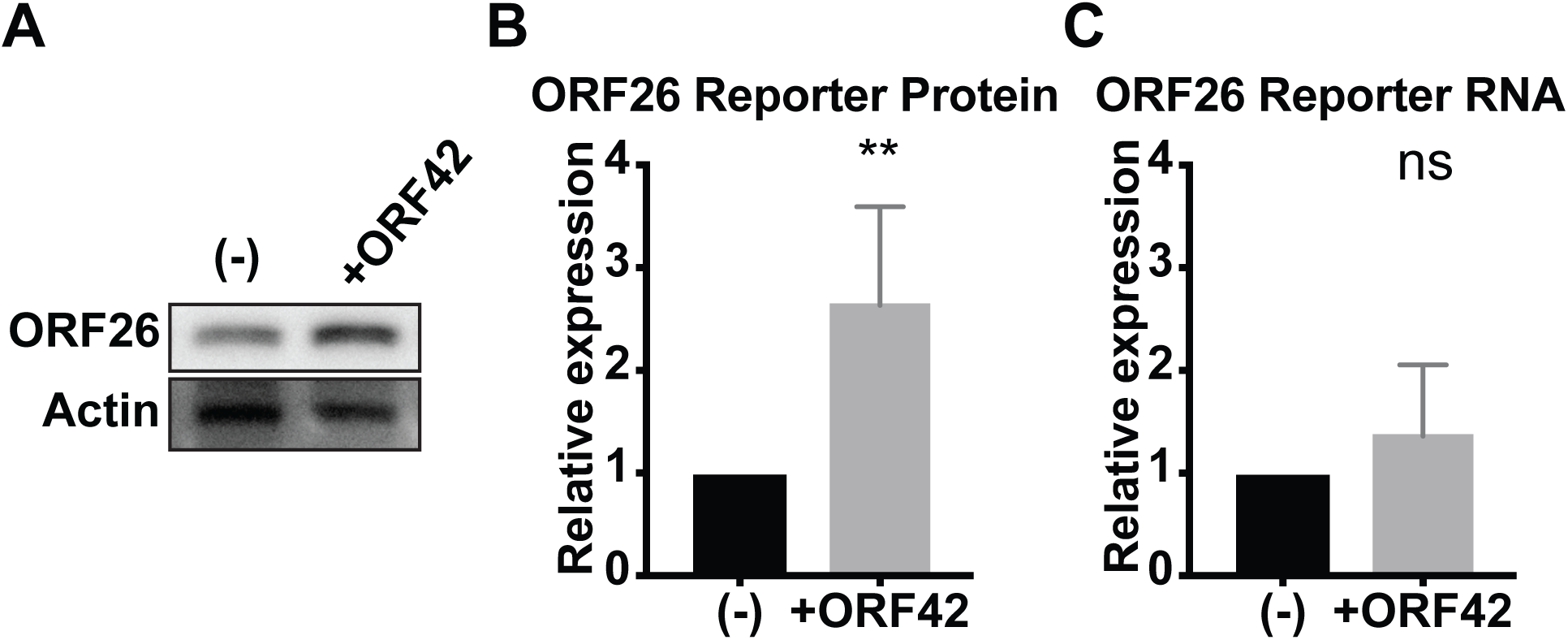
ORF42 increases levels of a co-transfected viral protein without altering mRNA levels. A construct expressing the ORF26 coding region and UTRs was co-transfected with either ORF42 (untagged) or empty vector. Protein and RNA were collected 24 hours later. (A-B) ORF26 protein was measured by western blot and normalized to actin. Quantification of several biological replicates is plotted in B. (C) ORF26 RNA was measured by qPCR and normalized to cellular 18S rRNA. N = 5; Ns, ** = *p* > 0.05 or < 0.01, respectively (Student’s t-test).

**Figure 4.**
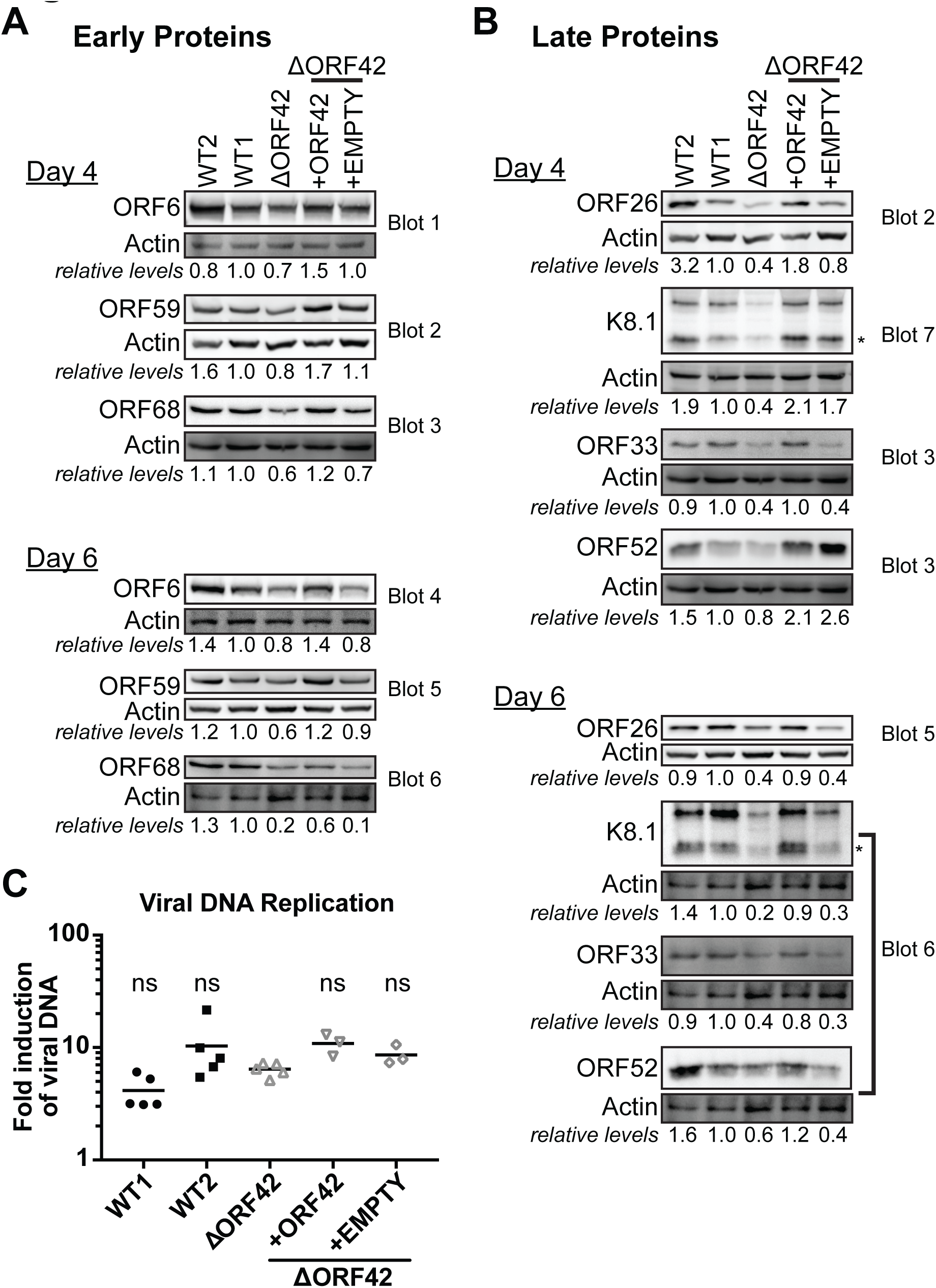
Loss of ORF42 impairs accumulation of viral proteins. The lytic cycle was induced in the KSHV-infected iSLK.RTA cells by addition of doxycycline (1 μg/ml) to the media. (A-B) Protein was collected four and six days post induction. Western blotting was used to measure the levels of the indicated early (A) and late (B) viral proteins. After quantitation, expression is reported normalized to actin and relative to WT1 line. All blots shown are from the same biological replicate and are representative of at least 3 biological replicates. Several proteins were measured on the same blot and normalized to the same actin blot, as indicated by the blot number. The band used to quantify K8.1 is marked by the asterisk. (C) Total DNA was collected from cells prior to induction and four days post induction. qPCR was used to measure copy numbers of the viral gene LANA and the cellular gene CCR5. The fold increase in DNA levels after induction was calculated after normalization to CCR5 levels. N ≥ 3. Ns = *p* > 0.05, relative to KSHV ΔORF42 (ANOVA followed by Dunnet’s multiple comparison test).

To determine whether ORF42 regulated the expression of viral proteins at the level of RNA or ; protein production, we measured RNA and protein in parallel for K8.1 and ORF26 (Fig. 5). i While we saw changes in both RNA and protein levels, the changes in protein levels seemed to l· precede those in RNA levels, at for at least K8.1. At day 4, the changes in RNA levels were not i statistically significant (Fig. 5A) and at day 6, the changes in RNA levels were smaller in i magnitude than those in protein levels (Fig. 5B). These results suggest a non-transcriptional role ‘ of ORF42. Also, because transcription of late genes is controlled by a complex of viral 1 transcription factors (5), changes in RNA levels could be indirectly caused by lower levels of these viral proteins. The lack of detectable ORF42 in the nuclear fraction (Fig. 2F) and the I increase in protein but not RNA of an ORF26 reporter upon ORF42 co-expression (Fig. 3) are also consistent with a predominantly non-transcriptional role for ORF42. Collectively, our ; results indicate that ORF42 enhances cellular and viral gene expression, at least in part by a post-transcriptional mechanism.

**Figure 5.**
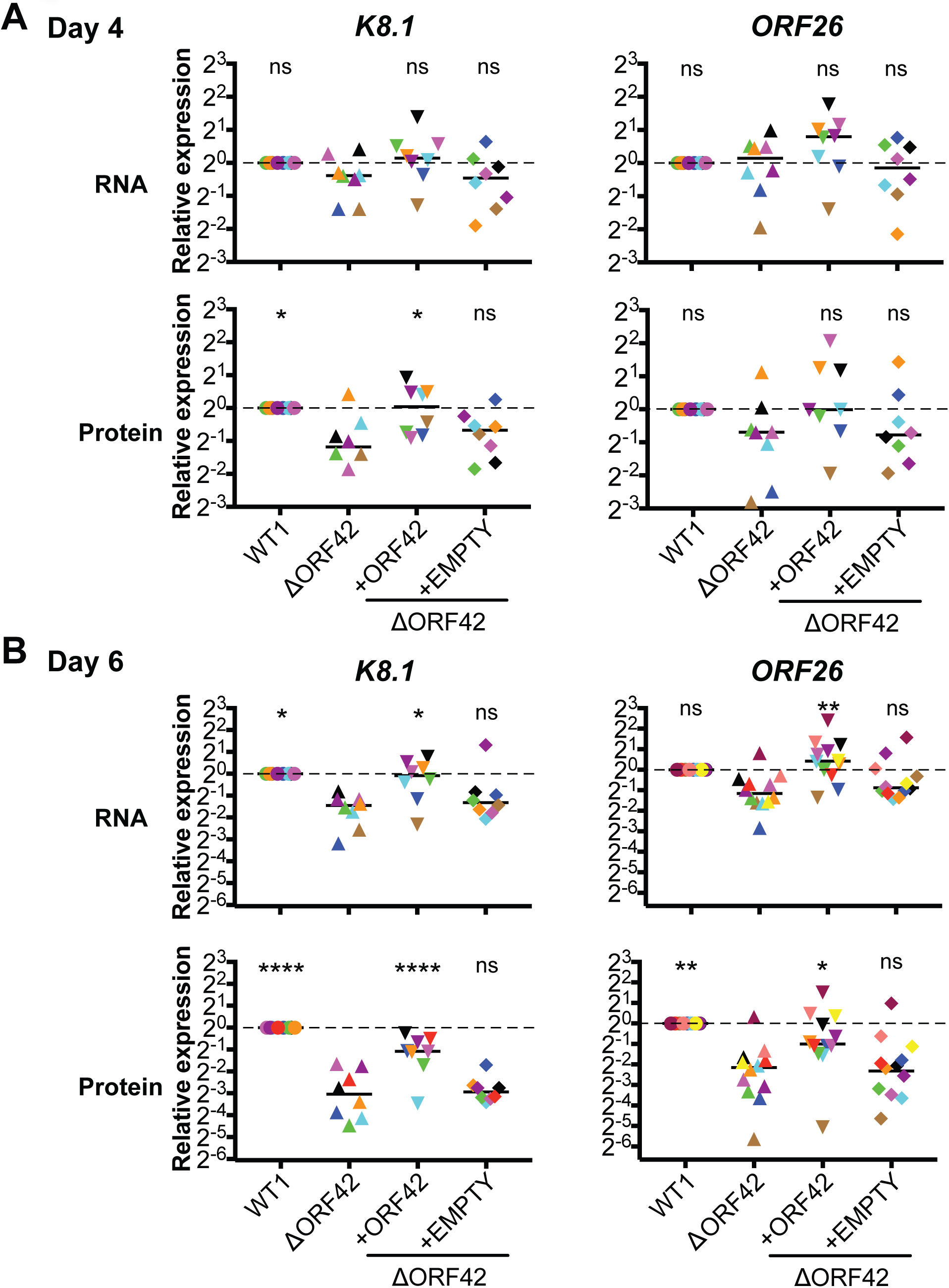
ORF42-dependent changes in viral protein levels precede changes in viral mRNA. The lytic cycle was induced in the KSHV-infected iSLK.RTA cells by addition of doxycycline (1 μg/ml) to the media. Protein and mRNA was collected (A) four and (B) six days post induction from the same biological replicate. Western blotting was used to measure the levels of the indicated viral proteins and expression is reported normalized to actin and relative to WT1 line. qPCR was used to measure the levels of the indicated viral mRNAs and expression is reported normalized to 18s and relative to WT1 line. N ≥ 8, colors represent individual replicates. Ns, *,**,**** = *p* > 0.05, or < 0.05, 0.01, 0.0001 respectively, relative to KSHV ΔORF42 (ANOVA followed by Dunnet’s multiple comparison test).

### ΔORF42-infected cells generate fewer viral particles

Although our results suggest that KSHV ORF42 can increase viral and host cell protein production, the main functions ascribed to its alpha- and beta-herpesviral homologs are in virion 1 formation and egress. In particular, some studies have suggested that ORF42 homologs directly regulate egress, because UL7/UL103 mutations reduce the levels of cell-free virus more I substantially than cell-associated virus (6, 7). In addition, the ORF42 homologs in alpha-and beta-herpesviruses are components of the tegument (7, 26–29), the layer of proteins between the ; herpesviral capsid and envelope. This localization is consistent with a role in virion formation, i since many proteins involved in virion formation and egress are also tegument components (30). l· However, whether ORF42 is a tegument protein has remained unclear, since only some of the i proteomic studies have detected ORF42 or its gamma-herpesvirus homologs in viral particles (26, 31-35). Using the KSHV-ORF42-Flag virus, we were able to determine that ORF42 was also incorporated in virions released into the supernatant. We detected ORF42-Flag in virions isolated by ultracentrifugation, as well as the capsid protein ORF26 (Fig 6A). It is unlikely that the ORF42 staining is due to cellular debris, because we did not detect the non-virion protein ORF59 in this pellet (Fig. 6A). We then tested how virion formation and egress were affected by ORF42 mutations. We found that the levels of virion-encapsulated DNA in the supernatant mirrored the relative changes in infectivity. KSHV ΔORF42-infected cells produced fewer viral particles than WT KSHV infected cells, and expression of ORF42 *in trans* rescued this defect (Fig. 6B). This result excludes the possibility that KSHV ΔORF42 virions are produced in equal numbers, but are less infectious. Moreover, we reasoned that if KSHV ΔORF42 had a defect in egress, as proposed for UL7 and UL103 (6, 7), there would be little or no ORF42-dependent difference in intracellular infectious virions. To test this idea, we isolated intracellular virus 6 days post induction of the lytic cycle using a freeze-thaw method (36) and measured infectious virus levels. We found that the relative levels of extracellular (Fig. 2C) and intracellular (Fig. 6C) infectious virions produced by WT and ΔORF42-infected cell lines as well as the ORF42 rescue lines were comparable (Fig. 6C). Collectively, these results indicate that ORF42 contributes to formation of viral particles, although we cannot discern whether this is because it is required for wild-type levels of virion components like ORF26 and K8.1 (Figs. 4, 5), or because ORF42 has dual functions in gene expression and virion assembly.

**Figure 6.**
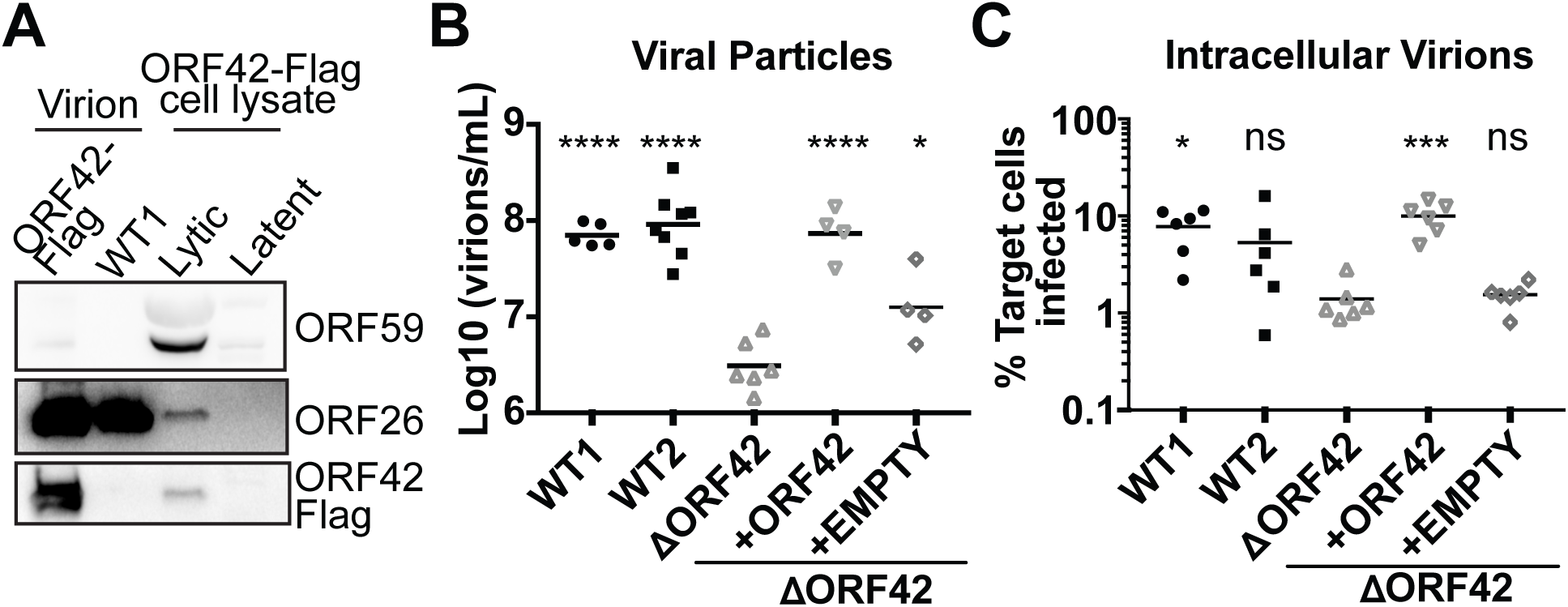
ΔORF42-infected cells are defective in virion formation. The lytic cycle was induced in KSHV-infected iSLK.RTA cells by addition of doxycycline (1 μg/ml) to the media. (A) Six days post induction, virions were isolated from the supernatant of cells infected with WT KSHV (“WT1”) or KSHV ORF42-Flag (“ORF42-Flag”) and lysed to collect protein. Lysates from latent and lytic KSHV ORF42-Flag-infected cells are shown as staining controls. Western blotting was used to detect ORF42-Flag, ORF59 (non-virion protein) and ORF26 (capsid protein). The blot is representative of 3 biological replicates. (B) qPCR was used to quantify levels of viral DNA in cell supernatant 6 days post induction, using primers against KSHV LANA. N ≥ 4. (C) Six days post induction, intracellular virions were isolated and used to infect HEK293T target cells. Infection was quantified by determining the percentage of GFP-positive target cells by flow cytometry. N = 6. For panels B and C, ns, * *** **** = *p* > 0.05, or < 0.05, 0.001, 0.0001 respectively, relative to KSHV ΔORF42 (ANOVA followed by Dunnet’s multiple comparison test).

## Discussion

We have identified a potential new function in cellular or viral gene regulation for four proteins of the gamma-herpesvirus KSHV: ORF22, ORF39, ORF42 and ORF55. Also, we uncovered evidence that five additional KSHV proteins that regulate viral gene expression (ORF18, ORF49, ORF59, K8α and K8β) could also have an effect on cellular gene regulation, since they up- or down-regulate expression of reporters that are controlled by non-KSHV regulatory regions. We further examined the role for one of the new candidate regulators, the previously uncharacterized protein ORF42, in gene regulation and in the viral replication cycle. We find that virus production is attenuated in cells infected with KSHV ΔORF42. Consistent with the identification of ORF42 from our screen for gene regulators, ORF42 up-regulates global protein synthesis during lytic infection and in transfected cells and is important for the accumulation of late viral proteins. ORF42 is predominantly detected in the cytoplasm and likely acts at a non-transcriptional stage of gene expression. Interestingly, we also found ORF42 in virions. This result, together with the fact that ORF42 homologs in alpha- and beta-herpesviruses function in virion formation, could point to a second role for ORF42 in virion assembly and/or a potential role for ORF42 in gene expression during *de novo* infections.

Although the homologs of ORF42 in other herpesviruses have generally been considered virion-assembly or egress proteins, we uncovered a role for ORF42 in gene expression. Indeed, during infection, nascent protein production and the accumulation of viral proteins are higher in the presence of ORF42. We do not yet know how ORF42 promotes protein accumulation, and whether it acts on select proteins or generally alters the translational output of the infected cell. The metabolic labeling results (Figs. 1C, 2G) may indicate that ORF42 affects global protein synthesis rates, but the small magnitude of the change suggests it has a somewhat selective effect. Also, based on our results, we conclude that ORF42 acts at a post-transcriptional stage of gene expression. The strongest evidence is that ORF42 overexpression up-regulates the protein levels of an ORF26 reporter without increasing its mRNA levels (Fig. 3). In addition, ORF42 is almost exclusively cytoplasmic, while nuclear localization may be expected for a direct effect on transcription (Fig. 2F). In the context of viral infection, ORF42 mutation decreases viral protein accumulation more dramatically than viral mRNA levels, although we did find that mRNA levels were also affected at day 6 post lytic induction (Fig. 5B). While ORF42 could affect multiple steps in gene expression, loss of ORF42 may result in decreased accumulation of virus-encoded transcription factors, which in turn could indirectly reduce viral transcription. In fact, mutations in KSHV ORF45 cause a similar and likely indirect reduction in viral mRNA levels (37). ORF45 regulates viral and host protein translation through activation of the ERK/RSK kinases and phosphorylation of the translation initiator factor eIF4B, but has no reported role in transcription (38). What would be the benefit of having KSHV-encoded post-transcriptional regulators, particularly one that boosts accumulation of late proteins like ORF42? Translation can be inhibited by stress and innate immune responses (39), so viral proteins that stimulate gene expression at a post-transcriptional stage could be important to maintain abundant viral protein production. It may also be advantageous to boost viral gene expression only in the later stages of the replication cycle, to promote virus formation without exposing the virus to recognition by the immune system too early in replication. Because ORF42 is itself expressed with late kinetics, we favor a model whereby ORF42 has a greater effect on late proteins simply because it is only expressed during late stages of viral replication. However, we cannot exclude the possibility that late protein translation requires specific cellular and/or viral factors like ORF42 that are dispensable for early protein expression, as is the case for late gene transcription.

While the reduction in virion and tegument protein levels in the absence of ORF42 could explain the reduced viral production, there is a discrepancy in the magnitude of the effect. The change in protein accumulation was less than ten-fold (Figs. 4, 5), while the defect in virus production was at least 50-fold (Fig. 2C). In addition, ORF42 was found in virions, presumably in the tegument (Fig. 6A), and components of the herpesvirus tegument are often required for proper virion formation, as shown in KSHV for ORF33, ORF38, ORF45, and ORF52 (40–42). These observations, together with the fact that ORF42 homologs in alpha- and beta-herpesviruses have been implicated in virion assembly and egress, may indicate that KSHV ORF42 has a dual function. Because of the reduced accumulation of virion and tegument proteins in cells infected with KSHV ΔORF42, we cannot accurately assess additional roles in virion assembly. We also cannot exclude that virion-associated ORF42 may have a role during *de novo* infection, similar to HSV-1 UL7 (10), for example in boosting early protein expression. However, we note that it is likely that few copies of ORF42 are present in viral particles, as most of the former mass spectrometry studies of virions of KSHV and the related gamma-herpesviruses Epstein Barr virus, MHV68 and rhesus monkey rhadinovirus did not detect ORF42 or its homologs in viral particles (26, 31-35)

Our characterization of one of the hits from our screen as a regulator of protein production, as well as the identification of several known regulators of viral gene expression, validates the results from the reporter screen, and suggests that the other novel candidates, ORF22, ORF39 and ORF55, could also have a role in gene regulation. Since our screen favors the identification of candidates that have broad effects on gene expression, we have likely missed regulators that have more specific effects on subsets of mRNAs or proteins. For example, ORF45 regulates translation (38), but was not identified in our screen. This is probably due to the fact that ORF45 promotes translation predominantly of mRNAs containing highly structured 5’ UTRs (43), which was not the case for our reporters. In general, the characterization of the other novel hits from our screen will improve our understanding of how KSHV controls both host and viral gene expression. In addition, further work on ORF42 will elucidate how this protein controls protein production during the late stages of the viral replication cycle.

## Materials and methods

### Cells

HEK293T cells were cultured at 37°C, 5% CO_2_ in Dulbecco’s modified Eagle’s medium (DMEM, Gibco/Thermo Fisher) supplemented with 10% fetal bovine serum (FBS, Hyclone). iSLK.RTA cells containing wild-type KSHV BAC16, BAC16 ΔORF42, or BAC16 ORF42-Flag were maintained in DMEM supplemented with 10% FBS and 400 μg/ml hygromycin (Enzo Lifesciences). KSHV-infected iSLK.RTA cells that also express a transgene were maintained in DMEM supplemented with 10% FBS, 400 μg/mL hygromycin (Enzo) and 100 μg/mL zeocin (Invivogen). Live cells were imaged using a Nikon eclipse TE2000-u.

### Plasmids

The library of Strep-tagged KSHV ORFs in the pCDNA4/TO-C-terminal-Strep backbone (12), pCDNA4/TO-ORF42-Flag (used for cloning ORF42), pCDNA4/TO-C-terminal-Flag, pJP1_Zeo and the TKp-luciferase construct (based on the pGL4.16 vector) were kind gifts by Dr. Britt Glaunsinger. pJP1_Zeo is a modified version of the pLJM-GFP vector in which the puromycin resistance gene was substituted with a zeocin resistance gene. The pBABE-d2GFP construct was created by cloning the d2GFP from the pd2eGFP-N1 construct (Clontech) into the pBABE vector. The hPGKp-DsRedDR construct was created by PCR amplifying the hPGK promoter from the pLJM-GFP vector (Addgene #19319) and cloning it into pCDNA3.1-DsRed-destabilized (a kind gift of Chris Sullivan), after excision of the CMV promoter with MluI and BamHI. To express ORF42 from a transgene, ORF42 was PCR amplified from pCDNA4/TO-ORF42-Flag with or without the Flag tag, and inserted into the AgeI and EcoRI restriction enzyme sites of the lentiviral vector pJP1_Zeo. To generate pCDNA4/TO-ORF42(untagged), ORF42 amplified from pCDNA4/TO-ORF42-Flag was re-inserted in the PmeI digested pCDNA4/TO-ORF42-Flag vector. pJP1-Flag was generated by excising the GFP gene from pJP1_Zeo, and replacing it with a 3xFlag tag using the restriction sites NheI and EcoRI. To generate the pCMV-ORF26 construct expressing the entire ORF26 mRNA, the ORF26 locus was amplified by PCR from BAC16 and inserted into the backbone of pd2eGFP-N1 digested with NheI and AflII (to remove GFP and the SV40 3’UTR). T4 DNA ligase-based cloning (New England Biolabs) or Gibson cloning (HiFi Assembly Mix, New England Biolabs) were used to generate the constructs.

### Screen

HEK293T cells were transfected with single Strep-tagged KSHV ORF constructs in the pCDNA4/TO backbone (12). They were also transfected with a vTK-promoter driven Firefly luciferase expression construct with a SV40 3’ UTR, or the pBABE-d2GFP construct, or a hPGKp-driven DsRed-DR construct with bovine growth hormone 3’ UTR. A ratio of 2:1 between the KSHV ORFs and the reporter construct was used. For the luciferase assays, cells were transfected in 96-well plates and luciferase activity was detected 24 hrs post transfection with Steady-Glo luciferase assay system (Promega). Each ORF was tested twice, and ORFs that increased or decreased luciferase levels were retested a third time. All ORFs that on average increased or decreased luciferase by ≥ 1.5-fold were tested for their effects on GFP expression at least three times. For the GFP assays, protein lysates were collected at 24 hrs post transfection and GFP levels were measured using western blotting, using tubulin as a loading control. A BioRad chemidoc system was used to acquire the Image and the ImageLab software to obtain the quantitations. ORFs that appeared to cause cell death were removed from further analysis. All ORFs that on average increased or decreased GFP protein by ≥ 1.5-fold were tested for their effects on DsRed-DR expression at least three times. For the DsRed assays, protein lysates were collected at 48 hrs post transfection and DsRed levels were measured using western blotting, using actin as a loading control. A Syngene system was used to acquire the Image and the Genetools software to obtain the quantitations. Expression of the ORFs was directly verified during the DsRed testing stage by western blotting, and ORFs that were not expressed well were eliminated from further analysis.

### BAC mutagenesis

KSHV BAC16 was kindly provided by Dr. Jae Jung and is described in Brulois *et al.* (23). Genetic manipulation of the BAC was performed in GS1783 *Escherichia coli,* which encodes the Red recombinase and the restriction enzyme I-Sce, using the protocol described by Tischer *et al.* (44). Briefly, a kanamycin-resistance (Kan) cassette flanked by I-Sce cleavage sites was amplified by PCR. The primers introduce sequences at each end of the Kan cassette that correspond to those at the desired insertion/mutation point. These sequences are also homologous to each other. For the ΔORF42 BAC, a serine-to-stop mutation in the amino acid 25 of ORF42 was inserted in the sequences flanking the Kan cassette. To construct the ORF42-Flag BAC, the 3’ end of ORF42 and the Flag sequence were fused to the 5’ end of the Kan cassette. The Flag sequence and the 5’ end of the overlapping portions of ORF42 and ORF41 were fused to the 3’ end of the Kan cassette. The PCR fragments were electroporated into BAC16-containing *E. coli* GS1783 cells. Red expression was induced by heat shocking the cells at 42°C in order to promote recombination of the homologous sequences. Recombinant clones were selected on Lysogeny broth (LB) plates containing 16 μg/ml chloramphenicol (Sigma-Aldrich) and 25 μg/ml kanamycin (Fisher Scientific) at 30°C. Insertion of the Kan cassette was confirmed by PCR. To remove the Kan cassette, I-Sce expression was induced by treatment with 1% L-arabinose, followed by induction of Red by heat shocking the cells at 42°C. Kanamycin-sensitive clones were isolated by replica-plating on kanamycin-chloramphenicol and chloramphenicol only arabinose-containing plates. DNA from multiple clones was purified. The integrity of the genome was verified by Restriction Fragment Length Polymorphism, and the mutations/insertions by sequencing the region of interest.

### Lentiviral transduction and transient transfection

To generate viral particles for lentiviral transduction, pJP1-based lentiviral expression plasmids were transfected with the packaging plasmids psPAX2 and pMD2 (Addgene #12260 and 12259) into HEK293T cells. Two days later, BAC16-containing iSLK.RTA cells were infected with filtered supernatant by centrifugation at 1,000 rpm for 1 hr in the presence of 8 μg/mL polybrene (Sigma-Aldrich). Transduced cells were selected using 250 μg/mL zeocin. To express proteins and RNAs with transient transfection, constructs were transfected into HEK293T cells using linear polyethylenimine (PEI, Fisher Scientific). 1.25 μg/mL of DNA and 3.75 μg/mL of PEI were used to transfect cells in 6-well plates.

### Virion measurements by flow cytometry and qPCR

The lytic cycle was induced in 1.02*10^6^ KSHV-infected iSLK.RTA cells by addition of doxycycline (1 μg/mL). Six days post-induction the supernatant was collected, and filtered to remove cells and other debris. HEK293T target cells were plated in 12-well plates at a density of 1.2*10^5^ cells/well and infected with virus-containing supernatant by centrifugation at 1,000 rpm for 1 hr in the presence of 8 μg/mL polybrene (Sigma-Aldrich). Three days post infection, cells were trypsinized and GFP-positive cells were counted by flow cytometry on a BD FACSCalibur at the Tufts Laser Cytometry core facility. FlowJo (BD Biosciences) was used to analyze the data. Titer was calculated as: (Number of target cells at infection) x (percent of GFP+ cells)/(volume of virus in mL). To isolate DNA from extracellular virions, 100 μl of supernatant were treated with 10 units of Turbo DNase (Ambion, Austin, TX) at 37°C for 1 hr to degrade extravirion DNA. The reaction was stopped by addition of EDTA (10 mM final concentration) and heating at 70°C for 15 min. 0.4 mg of proteinase K and 200 μl of AL lysis buffer (DNeasy kit, Qiagen) were added to lyse virions. The DNeasy kit was then used to isolate viral DNA following the manufacturer’s protocol. DNA levels were measured by qPCR using iTaq Universal SYBR reagent (Bio-rad), using primers against the LANA promoter (45). Levels were quantified against a standard curve generated by PCR amplification of each gene, in order to calculate the absolute number of DNA molecules, which was then used to calculate the number of particles per ml.

### Measurement of intracellular virions

Intracellular virions were isolated from infected cells using a protocol adapted from (36). 1.7*10^5^ KSHV-infected iSLK.RTA cells were and treated with 1 μg/ml doxycycline to induce the lytic cycle. Six days post-induction of the lytic cycle, the supernatant was removed and cells were washed with phosphate buffer saline (PBS) three times to remove extracellular virions. Cells underwent three freeze-thaw cycles in 2.5 ml DMEM supplemented with 10% FBS. Supernatant was then collected and filtered to remove cells and other debris. HEK293T target cells were plated in 24-well plates at a density of 6*10^4^ cells/well. For each viral strain, 2 mL of supernatant was added to one well in the presence of 8 μg/mL polybrene. Plates were centrifuged at 1000 rpm for 1 hour. Three days post infection, cells were trypsinized and GFP-positive cells were counted by flow cytometry on a BD FACSCalibur at the Tufts Laser Cytometry core facility. FlowJo (BD Biosciences) was used to analyze the data.

### Measurement of viral DNA replication

6.8*10^4^ KSHV-infected iSLK.RTA cells were plated and treated with 1 μg/ml doxycycline to induce the lytic cycle. Four days post-induction, DNA from cells and supernatant was collected and purified using the DNeasy kit according to the manufacturer’s protocol. Viral DNA was measured as described above for virion DNA. Levels of the cellular gene CCR5 (46) were measured to normalize by total DNA extracted. For each line, fold-induction of viral replication was calculated by dividing the normalized DNA levels in induced cells to those in uninduced KSHV-infected cells.

### Virion isolation

ORF42-Flag KSHV-infected iSLK.RTA cells were plated at a density of 3.06*10^6^ cells per dish on six 60-cm^2^ dishes. The lytic cycle was induced by addition of doxycycline (1 μg/mL). Six days post-induction the supernatant was collected, and filtered to remove cells and debris. The supernatant was centrifuged at 80,000 x *g* for 2 hours at 4°C through a 30% sucrose cushion in TNE buffer (50 mM Tris–HCl pH 7.4, 100 mM NaCl, 0.1 mM EDTA). The pelleted virions were resuspended in TNE buffer and underwent a second identical spin. The pelleted virions were then lysed and proteins were isolated using RIPA buffer (50 mM Tris–HCl pH 7.4, 1% NP-40, 0.5% Na-deoxycholate, 0.1% SDS, 150 mM NaCl, 2 mM EDTA).

### Protein isolation and western blotting

Cells were lysed and proteins were isolated using RIPA buffer. Lysates were cleared by centrifugation and total protein concentrations were quantified by Bradford assay (Bio-rad). Protein lysates were separated by SDS-PAGE and transferred to PVDF membranes. Membranes were blocked in 5% milk in PBST (PBS with 0.1% Tween-20 (Fisher)) and incubated with primary antibodies for 3 hours at room temperature or overnight at 4°C in PBST with 0.5% milk. Primary antibodies against the following proteins were used: KSHV ORF59, K8.1, ORF68 (all 1:10,000 dilution, gifts from Dr. Britt Glaunsinger (47, 48)) KSHV ORF26, ORF33 and ORF52 (1:1000 dilution with the exception of ORF26, used at 1:5, gifts from Dr. Fanxiu Zhu (42, 49)), KSHV ORF6 (1:500, gift from Dr. John Karijolich and Gary Hayward), Flag (1:500, Sigma), human β-actin (1:200, Santa Cruz Biotechnology), human β-tubulin (1:1,000, Sigma), GFP (1:1,000, Sigma), DsRed (1:200, Santa Cruz Biotechnology). Secondary anti-rabbit and mouse (Southern Biotech) or anti-goat (Southern Biotech or Santa Cruz Biotechnology) IgG antibodies conjugated to horseradish peroxidase were used at a dilution of 1:5000 in PBST with 0.5% milk. Membranes were stained using Pierce ECL western blotting substrate (Thermo Fisher) and imaged with a Syngene G:Box Chemi XT4 gel doc system. Images were quantified using GeneTools version 4.02.03.

### RNA isolation and RT-qPCR

Total RNA was extracted using a Quick-RNA MiniPrep kit (Zymo Research) and treated with Turbo DNase (Ambion/Thermo Fisher) before reverse transcription. For reporter mRNA measurements, cDNA was prepared using an iScript cDNA Synthesis Kit (Bio-Rad) per the manufacturer’s protocol. For viral mRNA measurements, cDNA was prepared using AMV RT (Promega) per the manufacturer’s protocol, using a cocktail of primers targeted at KSHV genes and 18S ribosomal RNA (rRNA), in order to obtain strand-specific measurements. In both cases, human 18S rRNA levels were used as an internal standard to calculate relative mRNA levels. Real-time qPCR was performed using iTaq Universal SYBR Green Supermix (Bio-Rad) in a CFX Connect Real-time PCR Detection System (Bio-Rad). No template and no RT controls were included in each replicate. The CFX manager software was used to analyze the data.

### Metabolic labeling

1.7*10^5^ cells KSHV-infected iSLK.RTA cells were plated and treated with 1 μg/ml doxycycline to induce the lytic cycle. HEK293T cells were transfected with either untagged KSHV ORF42 in the pCDNA4/TO backbone or empty pCDNA4/TO vector and pd2eGFP-N1 (to mark transfected cells). Five days post-induction or 24 hours after transfection, the media was replaced with methionine-free media for 1 hour, and then treated containing Click-iT® L-azidohomoalaine (AHA, final concentration 0.1 μM, Thermo Fisher). After 2-3 hours, cells were fixed in 4% paraformaldehyde, washed with PBS, blocked with 1X Permawash (BD Biosciences) in PBS for 30 minutes or overnight, and incubated in 1 μM Dylight 650-Phosphine in 1X Permawash in PBS for 2 hours to couple the fluorophore to AHA. After washing with PBS, AHA incorporation into proteins was measured by flow cytometry on a BD LSRII flow cytometer at the Tufts Laser Cytometry core facility. Flow cytometry data were analyzed using FlowJo version 10 to obtain the median fluorescence intensity (MFI) of cells. For transfected cells, only successfully transfected GFP-positive cells were used to calculate the MFI.

### Statistical analysis

All statistical analysis was performed using GraphPad Prism version 7.0b for Mac OS X, (GraphPad Software, La Jolla California USA). Statistical significance was determined by Student’s *t* test for individual comparisons or analysis of variance (ANOVA) followed by Dunnett’s multiple comparison test when multiple comparisons were required.

## Acknowledgements

We thank Dr. Glaunsinger for kindly providing the ORF library, several other constructs, and antibodies against KSHV proteins, and for help with the initial screening work. We also thank Drs. Zhu, Sullivan, Karijolich and Hayward for kindly providing antibodies and constructs. We thank Dr. Kristin Kotewicz, Erion Lipo, Ila Anand, Sarah Dupont and members of the Bohm and Moore lab at TUSM for technical advice and help. We thank Allen Parmelee and Stephen Kwok of the TUSM flow cytometry core for their technical assistance. We thank Drs. Munger and Moore and members of the Gaglia lab for critical reading of the manuscript. This work was supported by a Developmental Grant from the Lifespan/Tufts/Brown Center for AIDS Research (P30 AI042853) and an American Cancer Research Scholar award (131320-RSG-17-189-01-MPC) to MMG. HGR was supported by the Tufts Post Baccalaureate Research Education Program (R25 GM066567).

---

**Table 1.**
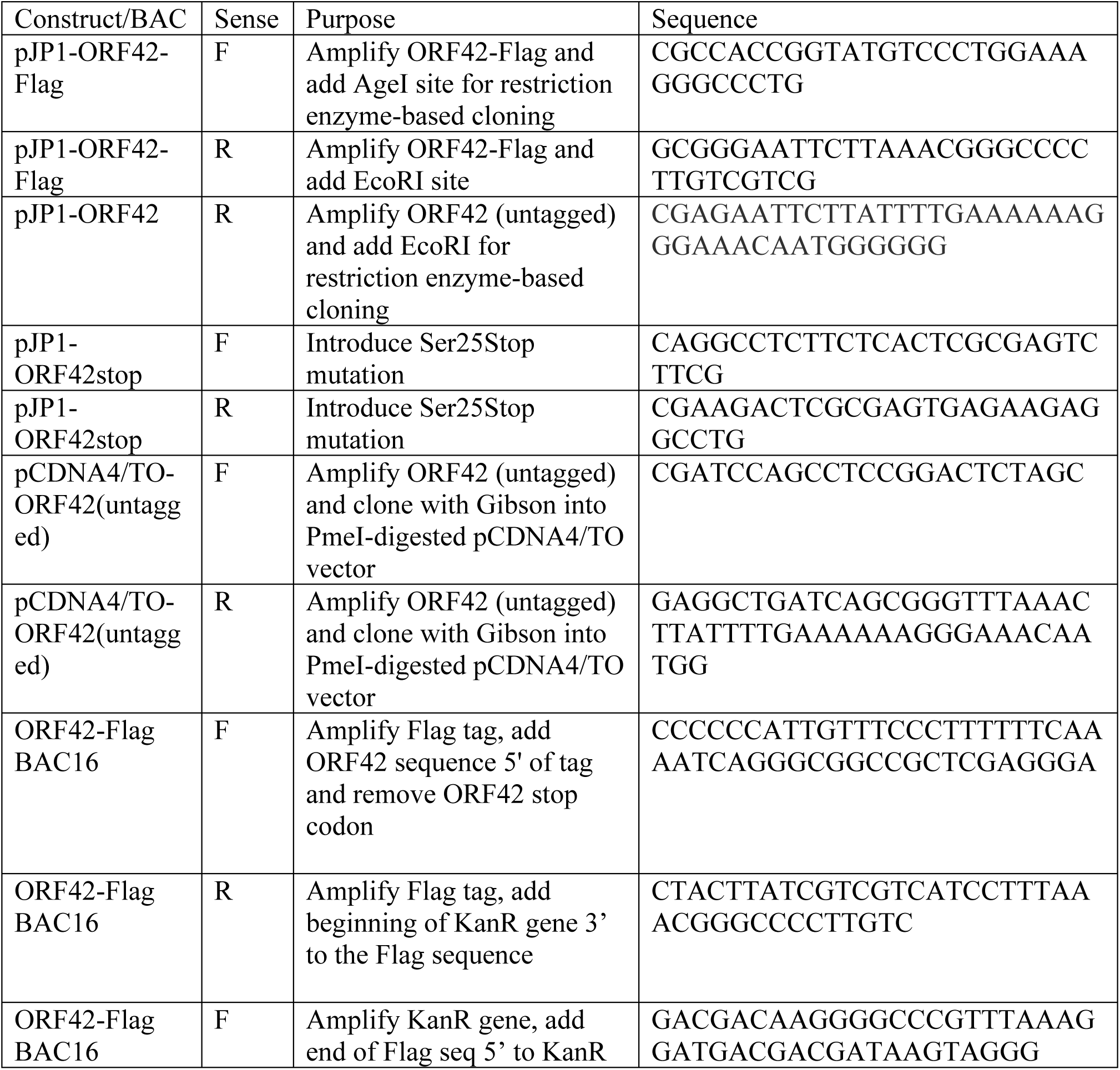
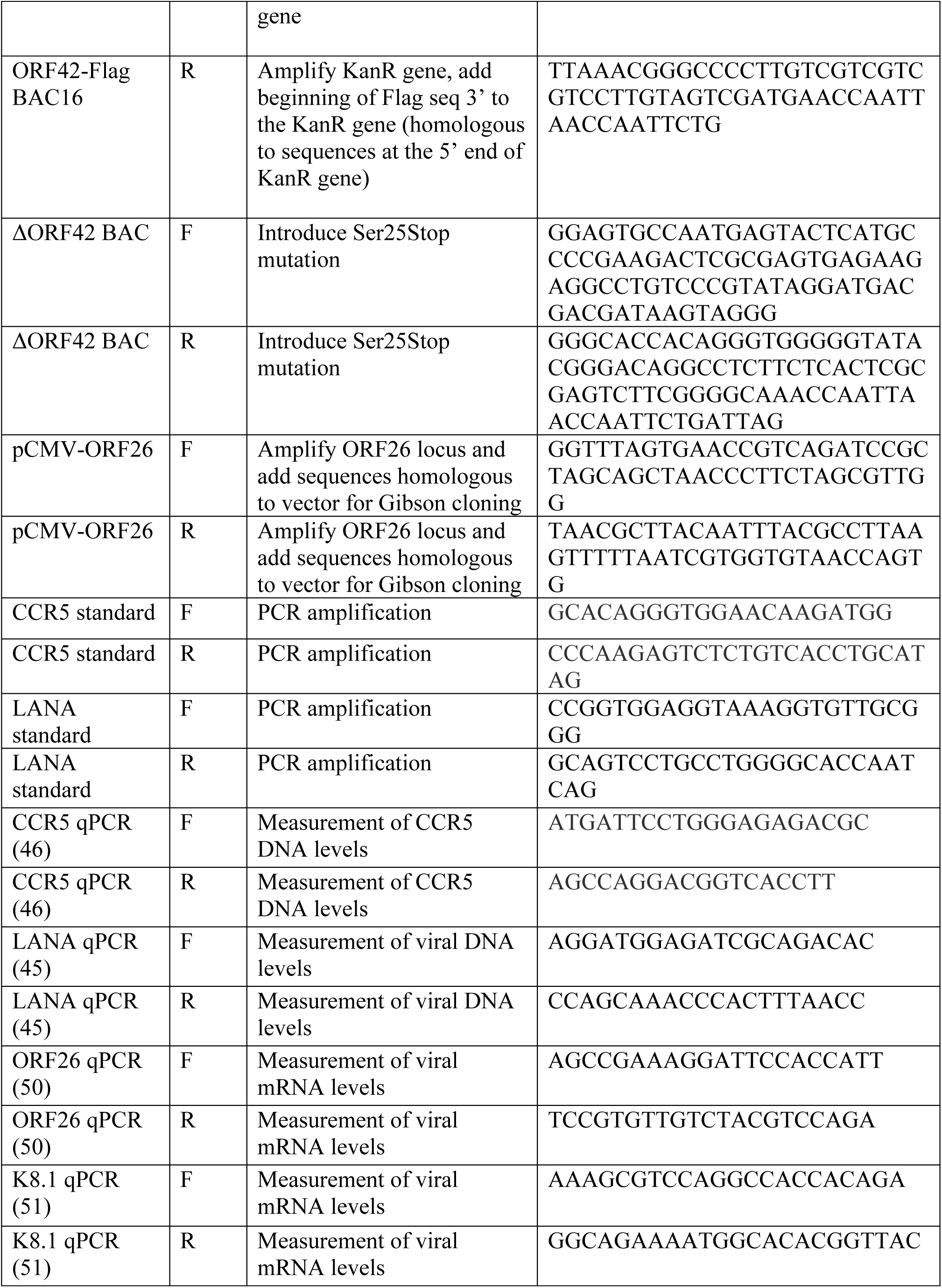
Primers.

